# Klf9 plays a critical role in GR –dependent metabolic adaptations in cardiomyocytes

**DOI:** 10.1101/2023.05.08.539871

**Authors:** Chandni Thakkar, Saleena Alikunju, Wajiha Rizvi, Ali Abbas, Danish Sayed

## Abstract

Glucocorticoids (GCs) through activation of the Glucocorticoid receptor (GR) play an essential role in cellular homeostasis during physiological variations and in response to stress. GC-GR signaling has been involved in regulating several cellular processes including metabolism, circadian rhythm and inflammation for diurnal adaptations. Our genomic GR binding (ChIP) and transcriptome (RNAseq) data from Dexamethasone (Dex) treatment in cardiomyocytes show an early (1hr) differential regulation of mostly transcription factors, followed by sequential change in downstream signaling pathways (6-24hr). Here, we examine the role of an early direct target of GR, Krüppel-like factor 9 (Klf9) in cardiomyocyte metabolic homeostasis. Our Klf9-ChIPseq identified 2150 genes with increase in promoter Klf9 binding in response to Dex. Functional annotation of these genes lists metabolic pathway on the top of KEGG pathway, along with genes regulating transcription and survival. Further, our transcriptome analysis of Dex treated cardiomyocytes with or without knockdown of Klf9 reveal differential regulation of 1777 genes, of which a reversal in expression is seen in 1640 (∼92%) genes with knockdown of Klf9 compared to Dex. Conversely, only 137 (∼8%) genes show further dysregulation in expression with siKLf9 as Dex treated cardiomyocytes. Gene ontology of these 1640 genes show metabolic genes on the top, including genes involved in glycolysis and oxidative phosphorylation. Expectedly, knockdown of Klf9 in cardiomyocytes inhibits Dex induced increase in glycolysis and spare respiratory capacity, as measured by glycolysis and mito stress tests, respectively. Thus, we conclude that cyclic, diurnal GC mediated GR activation, through Klf9 -dependent feedforward signaling plays a central role in maintaining cellular homeostasis through metabolic adaptations in quiescent and stressed cardiomyocytes.

## Introduction

Hypothalamus-Pituitary-Adrenal (HPA) axis regulates the cyclic release of Glucocorticoids (GC; cortisol in humans and corticosteroids in rodents) from adrenal cortex in response to physiological variations (circadian cycle, food), developmental cues, and stress conditions ^1^. Circulating GC can affect all tissues ^2, 3^, including cells of the heart and mediate their genomic actions by activation of cytosolic <u>n</u>uclear receptor subfamily <u>3</u>, group <u>C</u>, member <u>1</u> (Nr3c1) *aka* Glucocorticoid Receptor (GR), which translocate to the nucleus and associates with the chromatin at GR elements (GRE) ^4, 5^. Most of the GC actions are mediated through the genomic association of nuclear GR, with few non-genomic effects ^6, 7^. Glucocorticoids, through GR -dependent transcriptional targets regulate cellular rhythms, bioenergetics, growth and are potent anti-inflammatory agents ^8, 1, 9^.

GC function is required for postnatal cardiac growth and maturation ^10, 11^. Others and we have shown that Dexamethasone (Dex, synthetic GC with high selectivity for GR versus MR ^8^) mediated GR activation is sufficient for cardiac myocyte hypertrophy ^12, 13, 14^. We reported binding sites of GR across the cardiac genome, and identified early versus late, direct versus secondary transcriptional targets of activated GR in isolated cardiomyocytes treated with Dex ^14^. Our data shows that after 1hr of Dex stimulation 68 genes are identified as early direct targets of GR, which are associated with GR genomic binding and show differentially regulation at the transcript level. These genes are categorized as mostly transcriptional regulators by Ingenuity pathway (IPA) and DAVID analysis ^14^, suggesting that these factors could be serving as feedforward modulators of the GR signaling for an amplified, diverse downstream cellular response in cardiomyocytes.

In this study, we examine the role and contribution of Klf9, a member of the Krüppel-like factor (Klf) family in the mediating GR dependent metabolic adaptations in cardiomyocytes. Klf family of transcription factors consists of 17 known members that regulate various cellular processes, including differentiation, proliferation, apoptosis, metabolism with implications in development and diseases of cardiovascular, nervous, respiratory and immune systems ^15^. Our data suggests that GR associates with Klf gene structures and the differential expression of Klf genes is selective and maybe cell-type dependent. We show that Klf9, an early direct target of GR, in response to Dex regulates the expression of genes that are involved in glycolysis and mitochondrial oxidative phosphorylation, along with other pathways like nucleotide, purine and pyrimidine, and amino acid metabolism in cardiomyocytes. In addition, Klf9 is required for Dex dependent increase in glycolysis and spare respiratory capacity during mitochondrial stress, thus signifying an essential role of activated GR-Klf9 axis in cardiomyocyte metabolic adaptations under quiescent and stress conditions.

## Methods

### Animals

The work was done in accordance with US National Institute of Health *Guidelines for Care and Use of Laboratory Animals*. All animal protocols were approved by the Institutional Animal Care and Use Committee (IACUC) at the Rutgers New Jersey Medical School, Newark, NJ. Female Sprague Dawley rats with 1day old litter for isolation and culture of primary neonatal cardiomyocyte and C57/blk mice for postnatal heart samples were purchased from Charles River Laboratories.

### Neonatal Myocyte Culture and Treatments

Neonatal cardiomyocytes were cultured as previously described ^14^. Briefly, hearts from 1day old Sprague Dawley rats were isolated and dissociated. Two step enrichment of cardiac myocytes was performed using Percoll Gradient followed by differential pre-plating, to deplete any contaminating non-myocytes. Cells were plated in DMEM-F12 (High Glucose, with L-glutamine and HEPES) with 10% fetal bovine serum, after twenty-four hours medium was change to serum free medium. Cultured cardiomyocytes were treated with 100% methanol (solvent) or Dexamethasone (D1756, Sigma USA) 100nM for the indicated time periods. For gene expression manipulation, cardiomyocytes are infected with recombinant adenoviruses (constructed using protocol as previously described by Dr. Frank Graham ^16^) expressing shRNA against rat Klf9 (NM_057211) or shRNAs against luciferase gene (control) at the multiplicity of infection (MOI) of 10-20 particles /cell or as indicated.

### ChIP-Seq

Neonatal cardiac myocytes cultured from 40 hearts isolated from 1day old Sprague Dawley rat pups were treatment with methanol (control) or Dexamethasone (100nM) for 6hrs, cells were fixed, collected and sent to Active Motif for Klf9-ChIP-Seq. Klf9-CHIP-Seq was performed using anti-Klf9 antibody (Cat # A7196, ABclonal technologies) on 25μg of chromatin from control, Dex treated and Input samples by Illumina (NextSeq 500) sequencing.

### ChIP-Seq data analysis

Bioinformatics on the sequencing data generated was performed by Active motif, Inc., as done and described previously ^17, 14^. The Data explanation from Active Motif includes the following: *Sequence analysis* – 75-nt sequence generated by sequencing reads were mapped to the rn6 genome using BWA algorithm, and the information stored in BAM format. Only reads which align with no more than two mismatches and map uniquely are used in analysis. *Determination of Fragment Density* – Aligned reads (tags) are extended in silico at their 3’ ends to a length of 200bp using Active Motif software, corresponding to the to the average fragment length in the size selected library. To identify the density of fragments along the genome, the genome is divided into 32-nt bins and the number of fragments in each bin is determined, and stored in BigWig file and can be visualized on genome browsers. *Peak Finding* – Genomic regions with enrichments in tag numbers are termed ‘intervals’, and defined by the chromosome number, start and end coordinate. 15,828,819 normalized tags were used for peak calling using MACS 2.1.0 algorithm with default cutoff p value 1e-7. Peak filtering was performed by removing false ChIP-Seq peaks as defined within the ENCODE blacklist. *Merged region analysis* – To compare peak metrics between two samples, overlapping Intervals are grouped into ‘Merged Regions’, which are defined by start coordinate of the most upstream interval and end coordinate of the most downstream interval (=union of overlapping intervals; “merged peaks”). In locations where only one sample has an interval, this interval defines Merged Region. The use of Merged Regions is necessary because the locations and lengths of intervals are rarely exactly same when comparing different samples. *Annotations* – Intervals, Merged Regions, their genomic locations along with proximities to gene annotations and other genomic features are determined and presented in Excel spreadsheets. Average and peak fragment densities within Intervals and Active Regions are compiled. Data obtained was sorted based on the ratio of Dex vs. control and subsets were made for manuscript preparation with proper data representation and presentation.

The Merged Regions provides the peak metrics for samples in all peak regions and used to compare ChIP enrichments between samples in the assay. Pairwise comparisons were performed using the MaxTags followed by Log2ratio. To calculate the maximum number of tags (reads) multiply the average value of the tag per sample with the length of the regions and divide by constant 224. The 224 value represents the average length of the sequenced fragment in base pairs. As the signal is normalized in 32 bp bins along the genome, 7 bins of 32 bp length (224) is equivalent to the fragment length of 200 bp. The MaxTags filter out the regions with low signal values, which are expected to be more variable and therefore lead to unreliable, often exaggerated ratios. Cutoffs were set at 100 MaxTags for Klf9-ChIP-seq. The sorted genes were further separated based on Log2ration of ≤0.705

### RNA-Seq

Neonatal cardiac myocytes cultured similarly as for ChIP-Seq were treated with methanol (control) or Dexamethasone (100nM) for 6hrs or 12hrs. Cells were sent to Azenta Lifesciences for RNA isolation and RNAseq, followed by bioinformatics, which included data analysis. The data generation and analysis from Azenta includes the following - RNAseq library was prepared via rRNA depletion and Illumina HiSeq 2×150 bp sequencing was performed. Sequence reads were trimmed to remove possible adapter sequences and nucleotides with poor quality using Trimmomatic v.0.36. The trimmed reads were mapped to the Rattus norvegicus Rnor6.0 reference genome available on ENSEMBL using the STAR aligner v.2.5.2b. Alignment information for each read was stored in the BAM format. Only unique reads that fell within exon regions were counted and calculated by using featureCounts from the Subread package v.1.5.2. The hit counts were summarized and reported using the gene_id feature in the annotation file. Distribution of read counts in libraries were examined before and after normalization. The original read counts were normalized to adjust for various factors such as variations of sequencing yield between samples. These normalized read counts were used to accurately determine differentially expressed genes using DESeq2. Log2 fold change between the treatment groups was calculated and the Wald test was used to determine statistical significance (p*-*value). Adjusted p values were calculated using Benjamini-Hochberg procedure. Genes with an adjusted p<0.05 are reported as differentially expressed genes for the comparison between control and dexamethasone treated groups (Ad-siLUC vs ad-siLUC + Dex). Genes with an adjusted p<0.05 and log2FC of at least 0.5 were called out as differentially expressed genes for the comparison between dexamethasone treatments groups in the presence and absence of endogenous Klf9 knockdown (ad-siLUC + Dex vs ad-siKlf9+dex) Heatmap for the differentially regulated genes was generated using Heatmapper ^18^, with average linkage for clustering and Euclidean Distance Measurement Method.

### Genome Browsers

Integrated Genome Browser (IGB) ^19^ and Integrated Genomic Viewer (IGV) ^20^ was used for visualization of Klf9- and GR-ChIP-seq and RNAseq data, respectively.

### Functional annotation and GO terms

Functional annotation was performed using by Database for Annotation, Visualization and Integrated Discovery (DAVID) algorithm. Gene ontology terms and KEGG pathways are shown in Figures (top 10) and screenshots from DAVID showing the complete list is shown as supplementary figures.

### Seahorse Real-Time Cell Metabolic Assay

The mitochondrial respiration and glycolytic function parameters were measured using mitochondrial stress test and glycolysis stress test, respectively using a Seahorse XFe96 Extracellular Flux Analyzer (Agilent Technologies) according to manufacturer’s recommendations with modifications as per our experimental conditions. Briefly, neonatal cardiomyocytes were plated in 96-well Seahorse assay plates, and treated as described above. One hour prior to the measurements, the medium was replaced with Seahorse XF DMEM medium (pH 7.4) supplemented with 17.5 mM glucose and 100 uM palmitate for mitochondrial stress test and, 2mM glutamine and 100uM palmitate for glycolysis stress test. The cells were then incubated for 1 hour at 37°C without CO_2_. For mitochondrial stress test, after baseline oxygen consumption rate (OCR) measurements (basal respiration rate), 2 μM oligomycin, 1 μM FCCP, and 25 μM antimycin A plus 2.5 μM rotenone (A+R) were sequentially injected into each well. For glycolysis stress test, after baseline extracellular acidification rate (ECAR) measurements (non-glycolytic acidification), 10 mM glucose, 1 μM oligomycin, and 50 mM 2-deoxy-D-glucose (2DG) were sequentially injected into each well. The results were analyzed using Seahorse Wave 2.6 (Agilent). OCR measures the flux of oxygen and is proportional to mitochondrial respiration, while ECAR measures flux of protons and is proportional to glycolysis.

### Quantitative polymerase chain reaction (qPCR)

As described in ^21^. Briefly, total RNA extracted from hearts or isolated cardiomyocytes was reverse transcribed using High-capacity cDNA reverse transcription kit (ThermoFisher). TaqMan assays sets (Applied Biosciences) were used for qPCR using QuantStudio3 (Life technologies).

### Cellular fractionation and Western Blotting

As described previously ^22^. Complete list of the antibodies used with catalog numbers is provided in supplementary file.

### Statistics

ChIP-Seq statistics for Klf9 (provided by Active Motif) is shown in detailed ChIP-Seq analysis will be uploaded to GEO data repository. Calculation of significance between 2 groups was performed using an unpaired, 2-tailed Students *t* test (Excel software). All experiments were repeated three times, unless indicated and presented as average with SEM, p<0.05 was considered significant.

## Results

### Krüppel-like factor (Klf) family of transcription factors are early, direct binding targets of activated GR in cardiomyocytes, with a differential transcriptional regulation of 7 Klf genes

Our integrated genomic and transcriptome data (GEO accession # GSE114767) identified 68 genes that show genomic GR association and are differentially regulated after 1hr of Dex treatment in isolated neonatal ventricular cardiomyocytes ^14^. GR-ChIP-Seq data analysis revealed a GR genomic association with 12 of the 17 known members of the Klf family, some with multiple binding sites like Klf9, Klf13, and Klf15 suggesting that this family of transcription factors could be serving as early feedforward regulators of the GR signaling (**Fig 1A, 1B and 1C**). However, RNAseq results from isolated cardiomyocytes treated with Dex for increasing time points showed change in expression of just 7 Klf genes, including Klf2, Klf6, Klf9, Klf7, Klf10, Klf13 and Klf15 (**Fig 1A** and **S1A**). We observe an increase in expression of Klf9 and Klf15 as early as 1hr post Dex treatment that is maintained by 24hrs, while Klf10 expression decreases at 1hr, but reverts to endogenous levels by 6hrs (**Fig 1A** and **1D**). Klf2, Klf6 and Klf7 show an increase in transcription by 6-12hrs, and levels show a normalizing trend by 24hrs. Klf4, Klf3 and Klf16 with GR binding sites, and Klf11, Klf12 and Klf5 with no GR binding sites do not show significant difference in mRNA levels with any time points. Klf1 and Klf14 have GR binding, but the transcripts were too low and hence not included in further analysis (Fig S1B and S1C). On the hand, Klf8 and Klf17 are not associated with genomic GR and transcript did not sequence, suggesting very low or no expression in cardiomyocytes. These results identify selected Klfs as transcriptional targets of Glucocorticoid-GR signaling in cardiomyocytes. In addition, they also suggest that genomic GR binding may be independent of cell type and the change in mRNA abundance is regulated at different transcriptional or posttranscriptional level in a cell-type specific manner.

**Figure 1.**
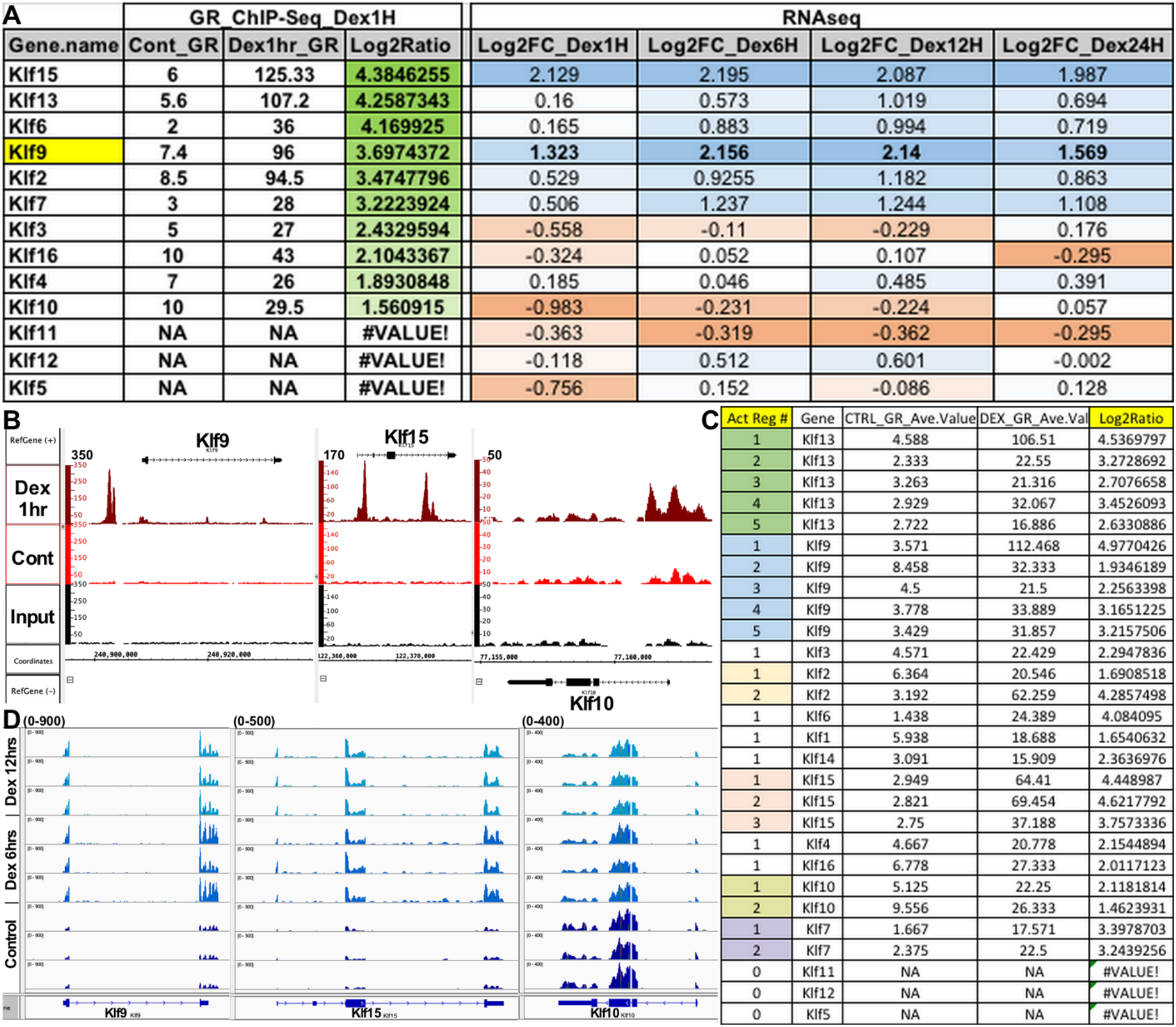
Activated GR regulates Klf family of transcription factors in cardiomyocytes. **A**. Table shows the genomic GR occupancy (GR-ChIP-seq) and transcript abundance (RNAseq) of Klf family from isolated neonatal cardiomyocytes treated with control (methanol) or Dex (100nM) for increasing time periods of 1hr, 6hr, 12hr or 24hrs, as indicated. The Klf genes shown are sorted based on differential GR occupancy (Log2FC, shades of green) with the corresponding changes in transcript levels (Log2FC, shades of red and blue). **B**. Screenshot of Integrated Genome Browser (IGB) showing the associated genomic GR occupancy on Klf9, Klf15 and Klf10 gene structures in control and Dex (100nM, 1hr) treated cardiomyocytes. The X axis represents chromosomal coordinates, reference sequence and gene structure, Y axis shows the peak value on the signal tracks of GR genomic occupancy in the samples (control, Dex and Input). **C**. Table shows the number of active regions (independent GR associating sites on Klf gene structures, along with the average peak density and Log2ratio of each region in control and Dex treated cardiomyocytes. **D**. Screenshot from Integrated Genome Viewer (IGV) showing the RNAseq data of representative Klf genes, Klf9, Klf15 and Klf10 in cardiomyocytes treated with control or Dex for 6hrs and 12hrs in triplicates. The X axis represents reference sequence and gene structure, while the Y axis shows values on the signal tracks. The value is constant across the samples for each gene (0-900 for Klf9, 0-500 for Klf15 and 0-400 for Klf10).

### Expression levels of Klf9, Klf15 and Klf10 correlate with GR activation status in neonate, young and adult mouse hearts

To examine the correlation of activated GR (nuclear vs cytoplasmic levels) with expression pattern of 3 early direct transcriptional Klf targets, we measured the transcript levels, protein abundance and cellular compartment localization of GR, Klf9, Klf15 and Klf10 in neonate (1d and 5d old), young (5wks old) and adult (12wks old) mouse hearts. Relative to the neonate hearts, which shows highest nuclear GR, we observe tight correlation between Klf expression pattern with nuclear GR abundance in these hearts. A reduction in nuclear GR (5d vs 1d old heart) is associated with a decrease in Klf9 and Klf15, and an increase in Klf10 levels. As the GR activation changes with age, Klf9 and Klf15 levels increase while Klf10 levels normalize to as in neonate hearts (**Fig 2A, 2C, 2D** and **2E**). Interestingly, the transcript abundance of GR does not change in these hearts, but protein expression decreases in the adult versus neonate hearts (**Fig 2A, 2C** and **2E**). Switch in myosin heavy chain (Myh) 7 to Myh6 expression confirms the postnatal development of these mouse hearts. Of the two additional GR targets in cardiomyocytes used for validation, Arrdc3 and Hes1 ^14^, Hes1 expression follows a similar pattern as Klf10 (**Fig 2B**). These results further validate Klf9, Klf15 and Klf10 as targets of activated GR, and suggest that Klf9, Klf15 and Klf10 act as essential early modulators of downstream GR signaling in heart.

**Figure 2.**
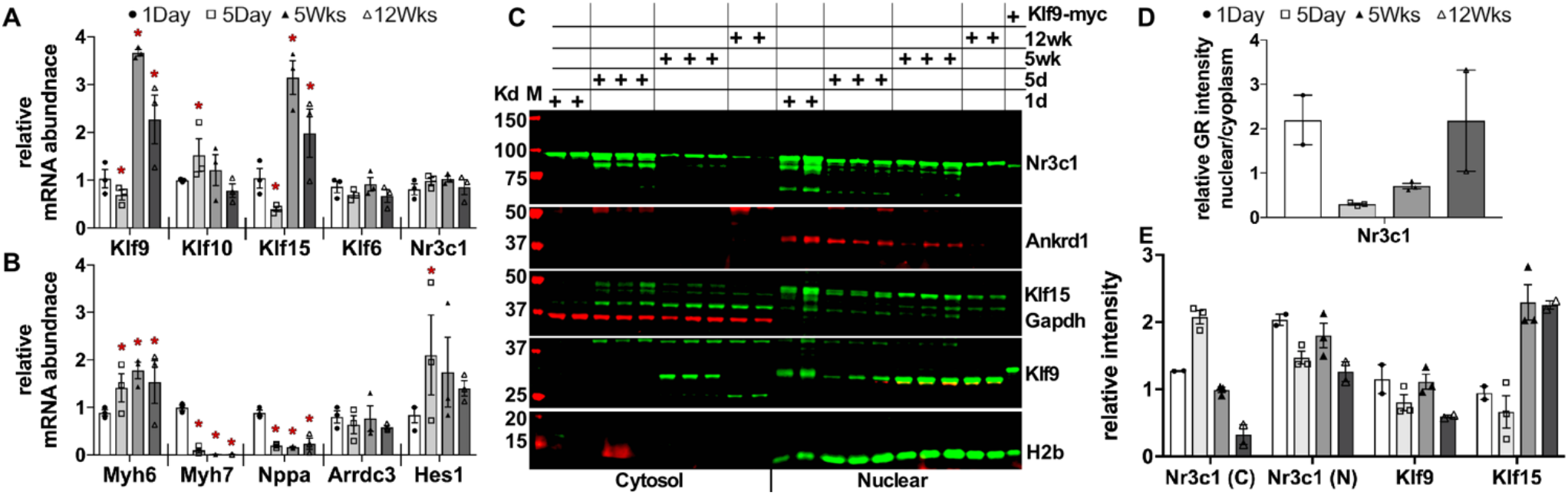
Nuclear GR expression correlates with differential Klf9, Klf15 and Klf10 expression with postnatal cardiac development. **A** and **B**. Graph represents qPCR data showing differential mRNA abundance of the listed genes from 5day, 5weeks and 12weeks relative to 1day old mouse hearts. Error bars represents SEM, * is p-value<0.05 compared to 1day old neonatal hearts, n=3. **C**. Western Blot showing cytoplasmic and nuclear fractions of heart tissue from mice of the indicated age. Gapdh and H2b shown as loading controls and for compartment specificity. n = 2-3. **D**. Graph represents the ratio of nuclear versus cytoplasmic GR signal intensity normalized to respective compartment controls (Gapdh or H2b), as measured from 2C using Image J, between the samples for postnatal hearts. **E**. Graph represents signal intensity of cytosolic and nuclear GR (Nr3c1), Klf9 and Klf15, normalized to their respective compartment controls (Gapdh or H2b) as measured from 2C using image J. Error bars represents SEM, n=2-3, as indicated with the data points.

### Klf9 mediates GR -dependent metabolic adaptations in cardiomyocytes

Of the three Klf (Klf9, Klf15 and Klf10) genes showing differential regulation with 1hr of Dex, we investigated the role of Klf9 in mediating the downstream effects of GR signaling. To examine the transcriptional targets of Klf9 in response to Dex, we performed high resolution illumina sequencing after chromatin immunoprecipitation with anti-Klf9 antibody (Klf9-ChIP-seq) in cardiomyocytes treated with methanol (control) or Dex (100nM) for 6hrs. As expected, we see an increase in genomic Klf9 occupancy with Dex treatment, with majority of the peaks at the promoter proximal and intronic regions. The data also shows that in contrast to GR occupancy, Klf9 associates with the cardiac genome even in control cardiomyocytes (**Fig 3A, 3B** and **S2**). In Fig 3C, we show the motif sequence identified by HOMER motif analysis of the top 2500 peaks (**Fig 3C**), which are enriched from HOMER’s internal motif database (**Fig S3**). Of the 15,558 merged regions with peaks for Klf9, 5284 were unique only to Dex treatment while 2475 only in control cardiomyocytes and 7799 were common in both control and Dex treated cardiomyocytes (**Fig 3D**). We further sorted the genes based on the MaxTag (≥100) and Log2FC (≥0.705). Of the 2155 genes sorted, 2150 showed an increase in Klf9 occupancy with Dex, while 5 showed decrease in Klf9 binding compared to control. We performed gene ontology on the 2150 genes that displayed increase in Klf9 genomic binding with Dex using Database for Annotation, Visualization and Integrated Discovery (DAVID) algorithm 6.8 ^23, 24^. Kyoto Encyclopedia of Genes and Genome (KEGG) pathway ^25^ analysis listed genes of the metabolic pathways at the top followed by PI3K-Akt signaling and cellular senescence (**Fig 4A**). GOTERM (Gene ontology) on these genes identified genes involved in transcription, apoptosis, and aging (**Fig 4B**). Fig 4C shows Klf9 and GR occupancy on representative genes (for metabolic pathways) along with the Input signals in control and Dex treated cardiomyocytes, while the table shows the tabulated average Klf9 and GR tags for these genes (**Fig 4C** and **4D**).

**Figure 3.**
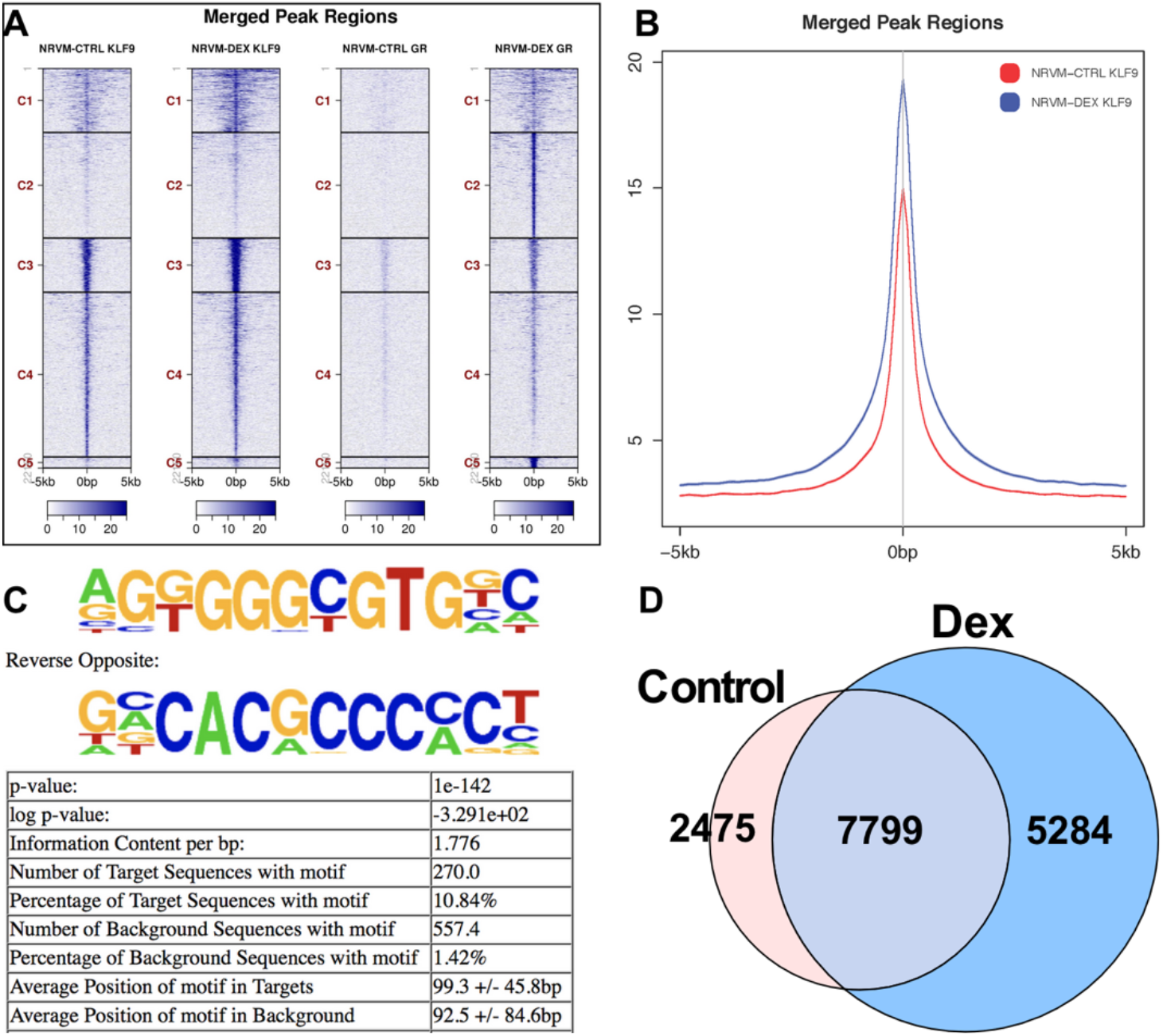
Dex stimulation increases genomic Klf9 association in cardiomyocytes. **A**. Heatmap showing tag distribution for Klf9 and GR across active regions (values in z-axis/color, active regions in y-axis) in control and Dex treated cardiomyocytes (6hrs for Klf9-ChIP-Seq and 1hr for GR-ChIP-Seq). The date is presented in 5 clusters (default), C1 to C5 and sorted. **B**. Average plots of Klf9 tag distribution of active regions (merged peak regions, including promoters and gene bodies) in control (red), Dex 6hrs treated (blue) cardiomyocytes. **C**. Enriched motif identified using Homer Motif analysis using the 200bp surrounding the summit of the top 2,500 peaks (based on MACS2 p-values). **D**. Venn diagram showing number of genes with Klf9 genomic association from Klf9-ChIP-Seq data in control and Dex (100nM for 6hrs) treated cardiomyocytes.

**Figure 4.**
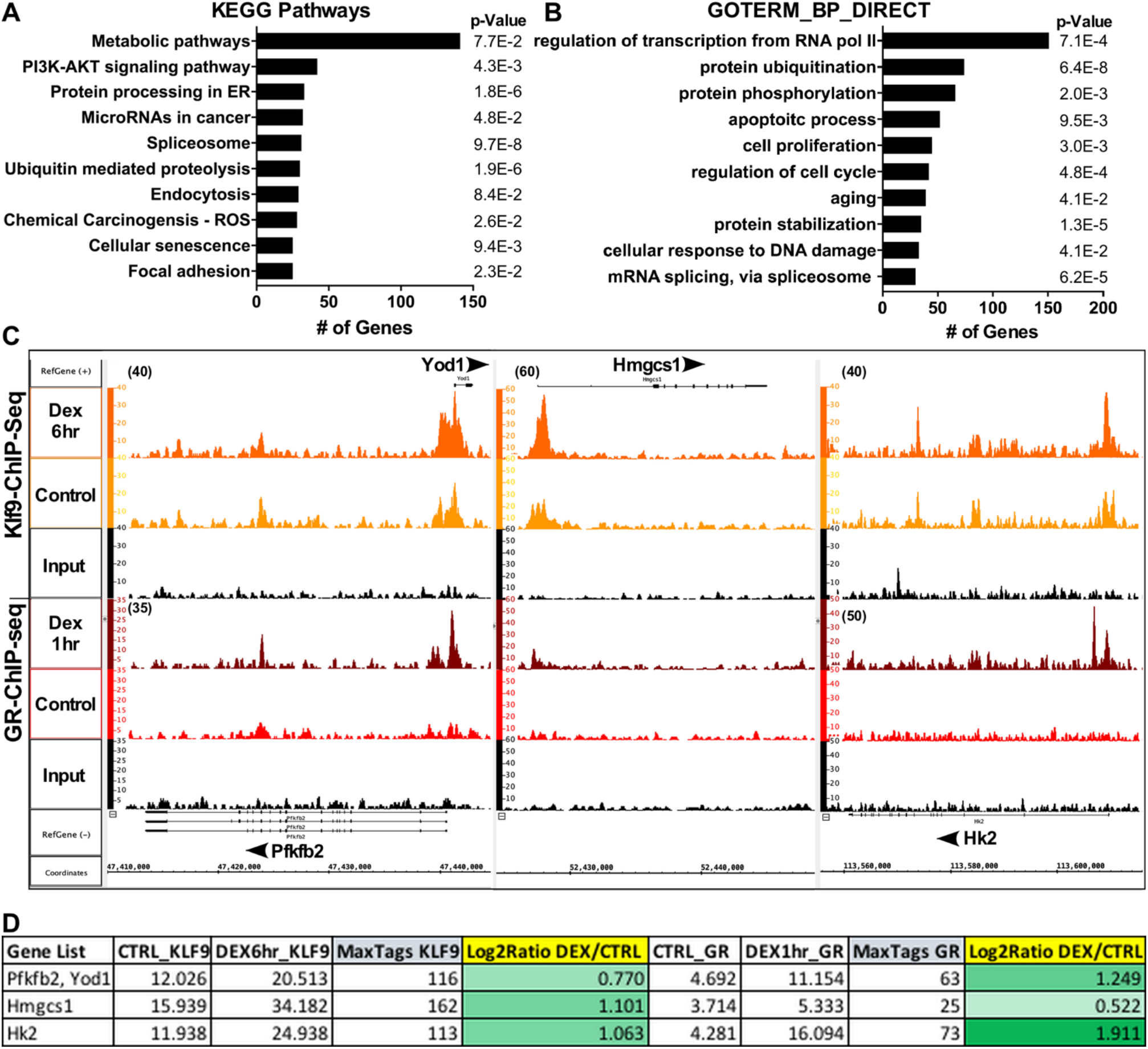
Klf9 associates with genomic regions of genes of metabolic pathways in response to Dex. **A** and **B**. Graphs lists the functional annotation and the number of genes that show Klf9 association in cardiomyocytes after Dex treatment. KEGG pathway and GOTERM analysis using DAVID was performed on 2150 genes which show increase in Klf9 genomic binding (Maxtags≥100, Log2FC ≥ 0.705). **C**. Screenshot of IGB showing Klf9 and GR occupancy on representative metabolic genes in cardiomyocytes treated with control or Dex (100nM) for the indicated time points. The X axis represents chromosomal coordinates, reference sequence and gene structure, Y axis shows the peak value on the signal tracks of Klf9 and GR genomic occupancy in the listed samples (control, Dex and Input). **D**. Table shows the average peak value of the Klf9 and GR on representative genes, along with the Max Tags and Log2FC.

Further, to examine the contribution of Klf9 in mediating Dex dependent change in gene expression, we performed total RNAseq in cardiomyocytes treated with Dex for 12hrs with or without knockdown of endogenous Klf9. Our data showed differential regulation of 7152 genes with Dex compared to control cardiomyocytes. Of these, 1777 genes show change in expression with knockdown of Klf9 (Ad-siKlf9) compared to Dex (Dex12h + Ad-siLUC) (**Fig 5A** and **5B**). Interestingly, 92.3% (1640) of the 1777 genes show reverse expression pattern with siKlf9 compared to siLUC in presence of Dex (12hrs), while transcript levels of 137 genes are further changed in the same direction as Dex (**Fig 5C**). KEGG pathway analysis of these 1640 genes shows metabolic pathways with 128 genes on the top of the list, followed by PI3K-Akt and MAPK signaling, and cellular senescence (**Fig 5D**). GOTERM categorized these genes as those involved in the regulation of transcription by RNA pol II, apoptosis, cell proliferation, aging and DNA damage (**Fig 5E**). Next, we sort the genes from the metabolic pathway identified by the KEGG pathway analysis, which categorizes 20 of the 128 genes as those involved in oxidative phosphorylation and diabetic cardiomyopathy. In addition, genes from purine and pyrimidine, amino acid, fatty acid and other metabolic pathways are modulated by GR-Klf9 signaling (**Fig 5F** and **5G**). In Fig 5H, we show the table listing the two essential genes from glycolysis pathway and the 20 genes from oxidative phosphorylation pathways that show reverse expression pattern with siKlf9 compared to control after Dex treatment for 12hrs in cardiomyocytes. These results from the ChIP and RNAseq data suggest that Klf9 is involved in regulating GR dependent metabolic homeostasis in cardiomyocytes.

**Figure 5.**
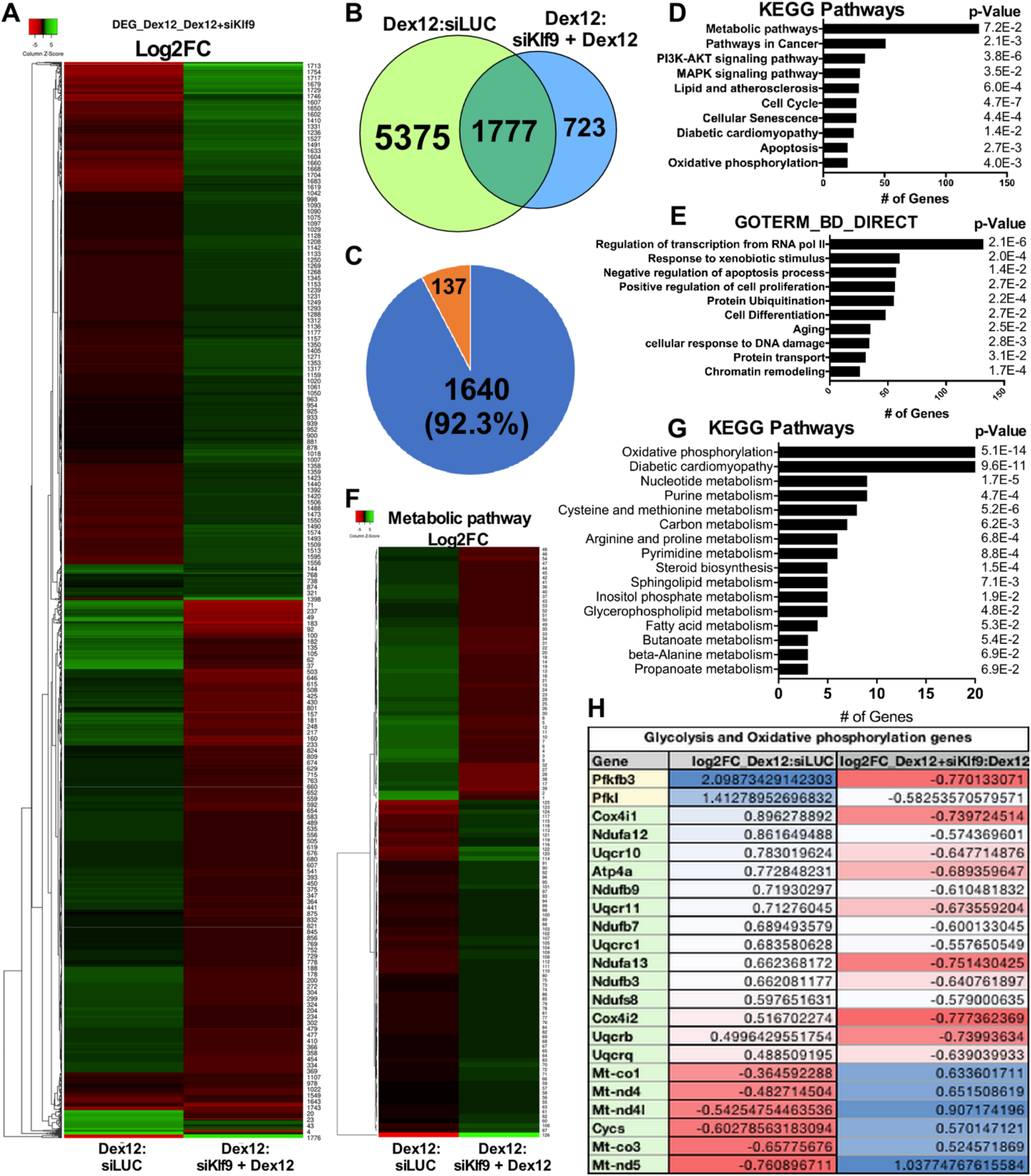
Klf9 is required for Dex dependent change in expression of genes involved in metabolic adaptations. **A**. Heatmap displays the 1777 genes that are differentially regulated in Dex (12hrs) treated cardiomyocytes in absence or presence of ad-siKlf9. Heatmap was generated using HeatMapper, with Average Linkage clustering method. RNAseq data included n=3 for each treatment. Gene list in excel sheet in attached as supplementary. **B**. Venn diagram showing number of genes that are differentially regulated in cardiomyocytes treated with Dex for 12hrs with or without Klf9 knockdown (ad-siKlf9). **C**. Pie chart showing the number of genes that show reverse (1640, 92.3%) in expression with Dex+siKlf9 versus Dex+siLUC (control), and genes (137, 7.7%) that show similar trend in expression with Dex+siKlf9 versus Dex+siLUC. **D** and **E**. Graphs lists the functional annotation with the number of related genes that show differential regulation with Dex with or without ad-siKlf9. KEGG pathway and GOTERM analysis using DAVID was performed on 1777 genes (Log2FC of 0.5). **F**. Heatmap displays the differentially regulated 128 genes associated with metabolic pathways identified by KEGG pathways analysis. Gene list in excel attached as supplementary. **G**. Graph shows the functional annotation and number of genes associated with different metabolic pathways from 128 genes from metabolic pathways identified by KEGG analysis. **H**. Table lists the genes, along with Log2FC that are involved in glycolysis and oxidative phosphorylation, and differentially regulated through GR-Klf9 axis.

### Klf9 is required for GR dependent increase in glycolysis and mitochondrial spare respiratory capacity in cardiomyocytes

We measured the baseline change in glycolysis and mitochondrial respiration in cardiomyocytes treated with Dex for increasing time periods. A significant increase in glycolysis after 1hr of Dex is observed, which further increases to maximum capacity at 6hr and 12hrs, before it starts normalizing by 24hrs (**Fig 6A, 6B and S4A**). Increase in glycolytic capacity is seen after 6hrs, while no significant changes are observed in glycolytic reserve between the control and Dex treated cardiomyocytes (Fig S4B). On the other hand, we did not observe a change in basal and maximum mitochondrial respiration with Dex, however, an increase is seen in the spare respiratory capacity with Dex treatments (**Fig 6C, 6D** and **S4B**). These results are suggestive of a significant role of Dex in metabolic homeostasis and adaptations in cardiomyocytes under quiescent and stressed conditions, respectively. Based on our genomic and transcriptome data, we predict that Klf9 is required for mediating the GR dependent metabolic alterations in cardiomyocytes. To that end, we performed Glycolysis stress and Mito stress tests with or without knockdown of Klf9 in the presence of Dex treatment for 12hrs. The results show that knockdown of Klf9 inhibited Dex dependent increase in glycolysis (**Fig 6E, 6F** and **S4D**) and mitochondrial spare respiratory capacity in cardiomyocytes (**Fig 6G, 6F** and **S4C**).

**Figure 6.**
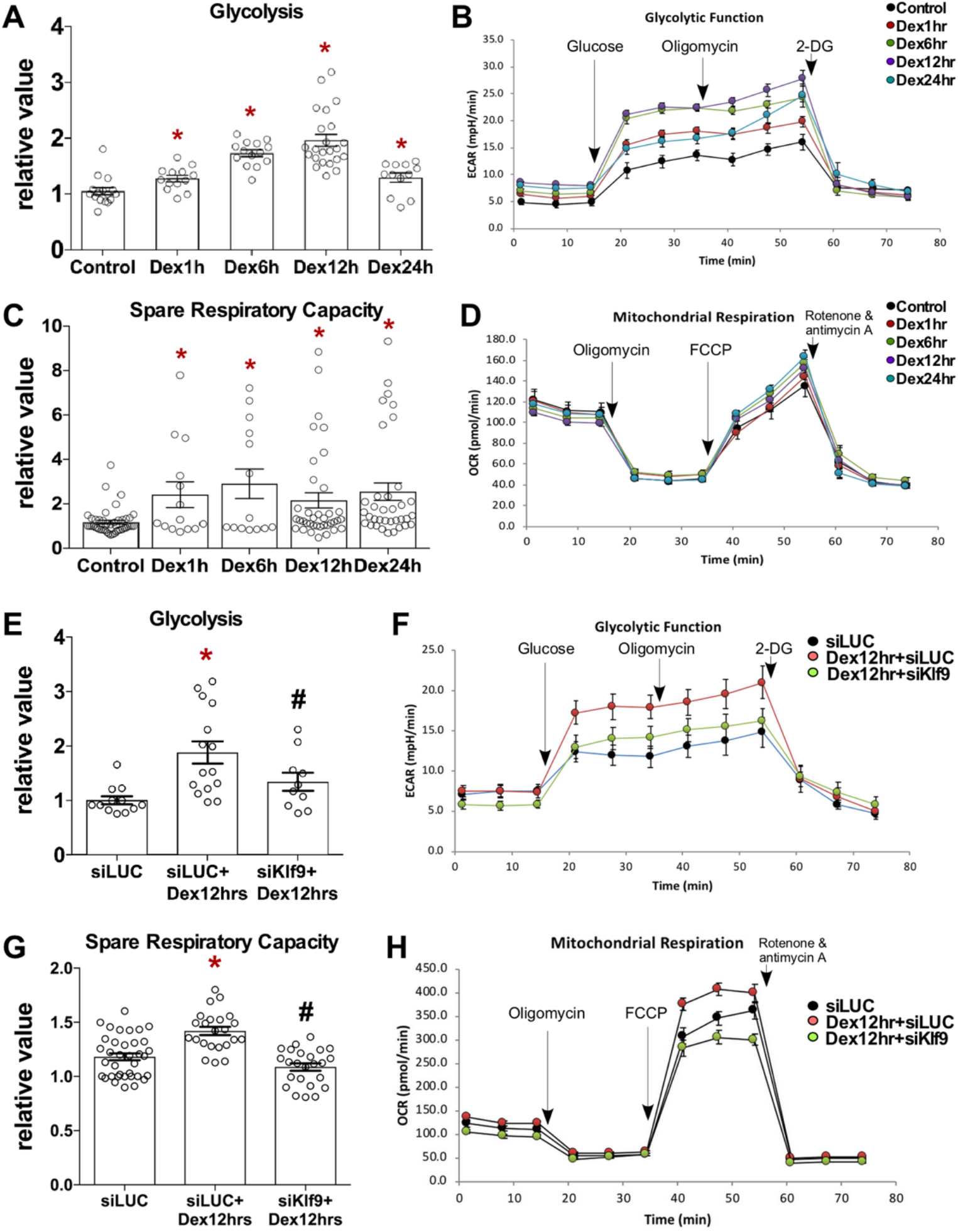
GR-Klf9 axis regulates metabolic adaptations in cardiomyocytes. **A**. Graph represents glycolysis in neonatal cardiomyocytes treated with Dex (100nM) for increasing time periods of 1hr, 6hr, 12hr and 24hrs. Each oval represents a data point from 3 independent cardiomyocyte cultures. Error bars represents SEM, p-value <0.05 compared to control. **B**. Representative graph of Glycolysis stress test, measuring the flux of protons, the extracellular acidification rate (ECAR) in neonatal cardiomyocytes treated with Dex for indicated time points. Glucose, Oligomycin and 2-DG were injected into the wells at the indicated time points during the assay to measure glycolysis, glycolytic capacity and glycolytic reserve. Error bars represents SEM. **C**. Graph represents spare respiratory capacity, as measured during the Mitochondrial stress assay in neonatal cardiomyocytes treated with Dex for increasing time points as indicated. Each oval represents a data point from 3 independent cardiomyocytes cultures. Error bars represents SEM, p-value <0.05 compared to control. **D**. Representative graph shows the Mitochondrial stress test, measuring flux of oxygen, the oxygen consumption rate (OCR) in neonatal cardiomyocytes treated with Dex for the indicated time periods. Oligomycin, FCCP and Rotenone + antimycin A were injected at the indicated time points during the assay to measure basal respiration, ATP production, proton leak, Maximal respiration and spare respiratory capacity. Error bars represents SEM. **E**. Graph represents glycolysis in neonatal cardiomyocytes treated with Dex (100nM) in absence or presence of Ad-siKlf9 or Ad-siLUC (control). Each oval represents a data point from 3 independent cardiomyocyte cultures. Error bars represents SEM, p-value <0.05 compared to control. **F**. Representative graph of Glycolysis stress test, measuring the ECAR in neonatal cardiomyocytes treated with Dex (100nM) in absence or presence of Ad-siKlf9 or Ad-siLUC (control). Glucose, Oligomycin and 2-DG were injected into the wells at the indicated time points during the assay to measure glycolysis, glycolytic capacity, and glycolytic reserve. Error bars represents SEM. **G**. Graph represents spare respiratory capacity, as measured during the Mitochondrial stress assay in neonatal cardiomyocytes treated with Dex (100nM) in absence or presence of Ad-siKlf9 or Ad-siLUC (control), as indicated. Each oval represents a data point from 3 independent cardiomyocyte cultures. Error bars represents SEM, p-value <0.05 compared to control. **H**. Representative graph shows the Mitochondrial stress test, measuring OCR in neonatal cardiomyocytes treated with Dex (100nM) in absence or presence of Ad-siKlf9 or Ad-siLUC (control). Oligomycin, FCCP and Rotenone + antimycin A were injected at the indicated time points during the assay to measure basal respiration, ATP production, proton leak, Maximal respiration and spare respiratory capacity. Error bars represents SEM.

Based on our genome wide occupancy data with Klf9 in cardiomyocytes in response to Dex treatment (Klf9-ChIP-Seq) and transcriptome data with knockdown of endogenous Klf9 (RNAseq) we conclude that Klf9 plays an essential role in mediating the downstream metabolic actions of GR signaling in cardiomyocytes. Dex dependent increase in Klf9 expression in cardiomyocytes is required for adaptive increase in glycolysis and mitochondrial spare respiratory capacity to maintain metabolic homeostasis during quiescent and stress conditions.

## Discussion

In this study, we extend from our previously reported genome-wide occupancy and transcriptome signature of activated GR in cardiomyocytes ^14^ to show that an increase in Klf9 expression is required for GR –dependent metabolic adaptations. Integrating of high throughput data sets with GR-ChIP and RNAseq, we show that nuclear GR associates with 12 out of 17 known Klf family members, but differential regulation of only 7 Klf genes is observed in cardiomyocytes with Dex treatment.

GR expression has been reported by 12wks and E9.5 -10.5d of gestation in human and mouse hearts, respectively, however, GR activation coincides with the surge in endogenous GC levels at late gestational period (E15.5-E17.5d in mice) and is required for organ maturation and postnatal survival ^26, 10^. Thus, as expected, global GR knockout presents with perinatal lethality in mice ^27^. Similarly, mouse fetus with conditional GR knockout under SM22a promoter develop cardiac dysfunction due to immature myofibrils and inhibition of cardiac-enriched contractile and metabolic genes ^10, 11^. On the other hand, αMHC-Cre induced GRKO (cGRKO), i.e., mice with postnatal GR knockout show spontaneous hypertrophy by 3mths, which progress into failure and death by 5mths. In addition to the increase in genes involved in mediating inflammation, there was inhibition of calcium handling genes and transcriptional regulators like Klf15 ^28^. These studies show the critical role of GR activation during postnatal cardiac development. Our data from postnatal developing hearts shows tight correlation between nuclear GR, Klf9 and Klf15 expression pattern, suggesting that GR-Klf axis might be playing a critical role in physiological cellular adaptations, as the neonate grows into an adult.

GC dependent GR activation/inactivation cycle in adult hearts leads to changes in transcriptome for diurnal adaptations. However, with extreme exercise or pressure overload in mice, there is decrease in cardiac GR expression, which results in altered transcription of GR -dependent genes that precipitates ventricular remodeling and dysfunction ^29, 30^. Interestingly, it has also been shown that ventricular dysfunction observed in mice with adrenalectomy could be partially reversed by supplementing exogenous corticosterone ^31^, suggesting that cardiac GR signaling is critical for maintaining quiescent cellular homeostasis, and hampered GR signaling could be contributing to cardiac pathogenesis. Others and we have shown that Dex mediated GR activation is sufficient for cardiac myocyte hypertrophy ^12, 13, 14^. Using Dex (highly selective for GR vs. MR ^8^) in isolated cardiomyocytes we reported genomic binding sites of GR ^14^, one of the first comprehensive GR association study in cardiomyocytes. Interestingly, 12 Klf family members were observed as potential GR targets, however, integrating the transcriptome data showed differential expression of only 3 Klf genes (Klf9, Klf10 and Klf15) at 1hr post Dex and 6 (Klf9, Klf7, Klf6, Klf2, Klf13 and Klf15) at later time periods up to 24hrs. These data suggest that the 3 Klf genes (9, 10 and 15) could be serving as early mediators, and as feedforward transcriptional regulators for an enhanced, amplified response of GR signaling. Further, our data also suggests that GR genomic association could be independent of cell type, however, the change in gene transcription is cell type specific, as seen with differential regulation of selected Klf genes with respect to GR association.

Klf family are zinc finger proteins that function as transcription regulators and can function as activators or repressors. With a conserved C-terminal DNA binding domain, Klfs can have overlapping transcriptional targets, however, the regulatory N-terminal domain controls association with other factors that can have different effects on target gene expression. Based on structural similarities and association with corepressor Sin3A, both Klf9 and Klf10 have been predicted to function as transcriptional repressors, while no distinct binding partners have been identified for Klf15 ^15^. Klf15, with high expression in heart is anti-hypertrophic and anti-fibrotic ^32, 33^. On the other hand, genomic Klf10 knockout mice develop spontaneous cardiac hypertrophy and fibrosis ^34, 35^. Klf10 has been shown to regulate T-cell activation ^36^, inhibit cancer cell proliferation ^37^ and can induce apoptosis ^38, 39, 40^. Recently Klf2, Klf9 and Klf15 were shown as direct GR targets in rat brain ^41^. With no significant reports of GR-Klf9 signaling in the heart, Klf9 has been shown to regulate hepatic glucose metabolism ^42^, repress inflammation and cytoskeletal remodeling in neurons ^43^, and as GR target in macrophages ^44^ and pulmonary epithelial cells ^45^. Our Klf9-ChIP-Seq data from cardiomyocytes treated with Dex for 6hrs, showing maximum increase in Klf9 expression in our time course study, identified genes involved in metabolic pathway on top of the list. Interestingly, aligning the Klf9-ChIP-Seq data (6hr Dex) with our previous GR-ChIP-seq data (1hr Dex) showed some overlapping targets, which could suggest that Klf9 could be acting as feedback regulator for these genes, thus, tightly controlling the expression levels in timed manner. Recently, Klf9 has been shown to regulate the expression of Fkbp5, a known target of activated GR, thus serving as downstream modulator of glucocorticoid sensitivity and metabolic homeostasis in zebrafish ^46^. However, in cardiomyocytes these observations need further confirmation and detailed functional evaluation for better understanding the complexities of GR-Klf9 signaling.

Heart is an omnivore and utilizes all sources for energy based on demand and availability. Cardiac myocytes can use lipids and carbohydrates as their energy source, and substrate utilization changes as the heart matures from fetal into the adult heart. High glycolytic activity in fetal cardiomyocytes switches after birth to oxidative phosphorylation (reviewed in ^47^), and these metabolic adaptations in postnatal hearts are dictated by the increase in ATP demands and consumption. Although, these changes at birth overlap with the surge in GR activity, there are no reports that have established direct correlation with the metabolic adaptations and GR signaling. Our transcriptome and functional data in neonatal cardiomyocytes show that Dex mediated increase in Klf9 is required for increase in glycolysis and mitochondrial spare respiratory capacity. Spare reserve capacity, as calculated by the difference between the maximum and basal respiratory capacity, gives measure of the potential of the cell to increase mitochondrial bioenergetic capacity with increasing demands and during oxidative stress ^48, 49^. Interestingly, GR expression and activity decreases with pathological cardiac hypertrophy ^29, 30^, along with decrease in expression of Klf9 (data no shown), suggesting hampered GR-Klf9 signaling and altered cellular metabolic adaptations to stress could be contributing to the pathogenesis.

In summary, we present a critical role of Klf9 as feedforward transcription regulator of GR signaling in cardiomyocytes, which in response to postnatal development, physiological variations or oxidative stress maintains cellular metabolic homeostasis and adaptations, respectively.

## Supporting information

Table with genes for Heatmap Fig 5

## Author contributions

CT and SA performed experiments; WR a summer intern assisted in experiments; AB assisted in RNA seq data analysis; DS designed experiments, performed data analysis with figures and wrote the manuscript.

## Acknowledgements

We thank Dr. Maha Abdellatif for guidance and support, and Dr. Sadoshima, Chair of department of Cell Biology and Molecular Medicine, Rutgers New Jersey Medical School for support.

