## Supplementary Figures for "Klf9 plays a critical role in GR –dependent metabolic adaptations in cardiomyocytes"

### Figure S1

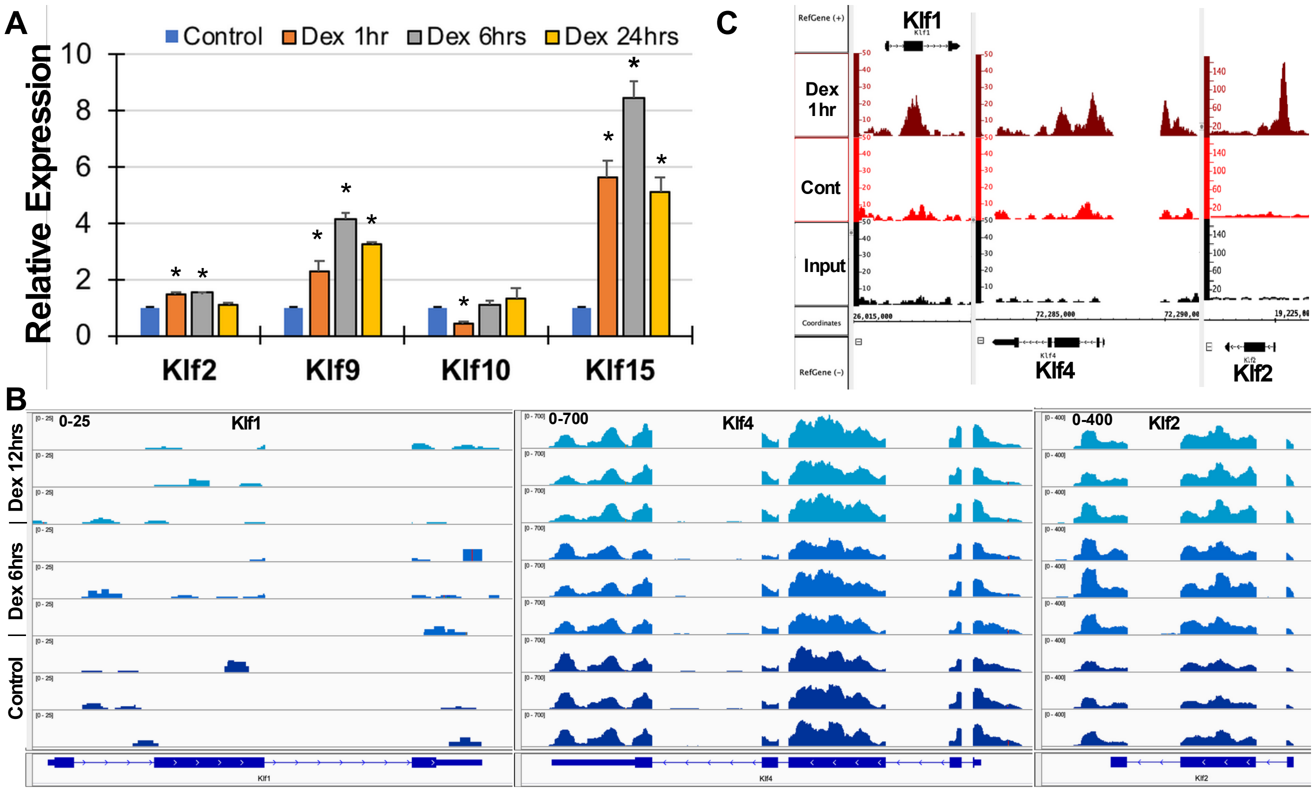


Figure S1. A. Graph represents qPCR data measuring the mRNA abundnace of the indicated genes with Dex treatment with increasing time periods relative to control. Error bars represents SEM, p value <0.05 compared to control, n=3. B. Screenshot of the IGV showing alignment of the RNAseq data of indicated genes from neonatal cardiomyocytes treated with Dex (100nM) for 6hrs or 12hrs on rat (rn6) reference genome. X axis represents ref sequnce and gene structure, while Y axis shows the values on the signal tracks. The value is kept constatnt across the samples for each gene. C. Screenshot of IGB showing the associated genomic GR ocupancy on Klf1, Klf4 and Klf2, as representative genes of the Klf family in cardiomyocytes treated with Dex or control for 1hr. X axis represents chromosomal coordinates, refernce sequnce and gene structure. Y axis shows the peak value on the signal tracks of GR in the control and Dex treated cardiomyocytes.

### Figure S2

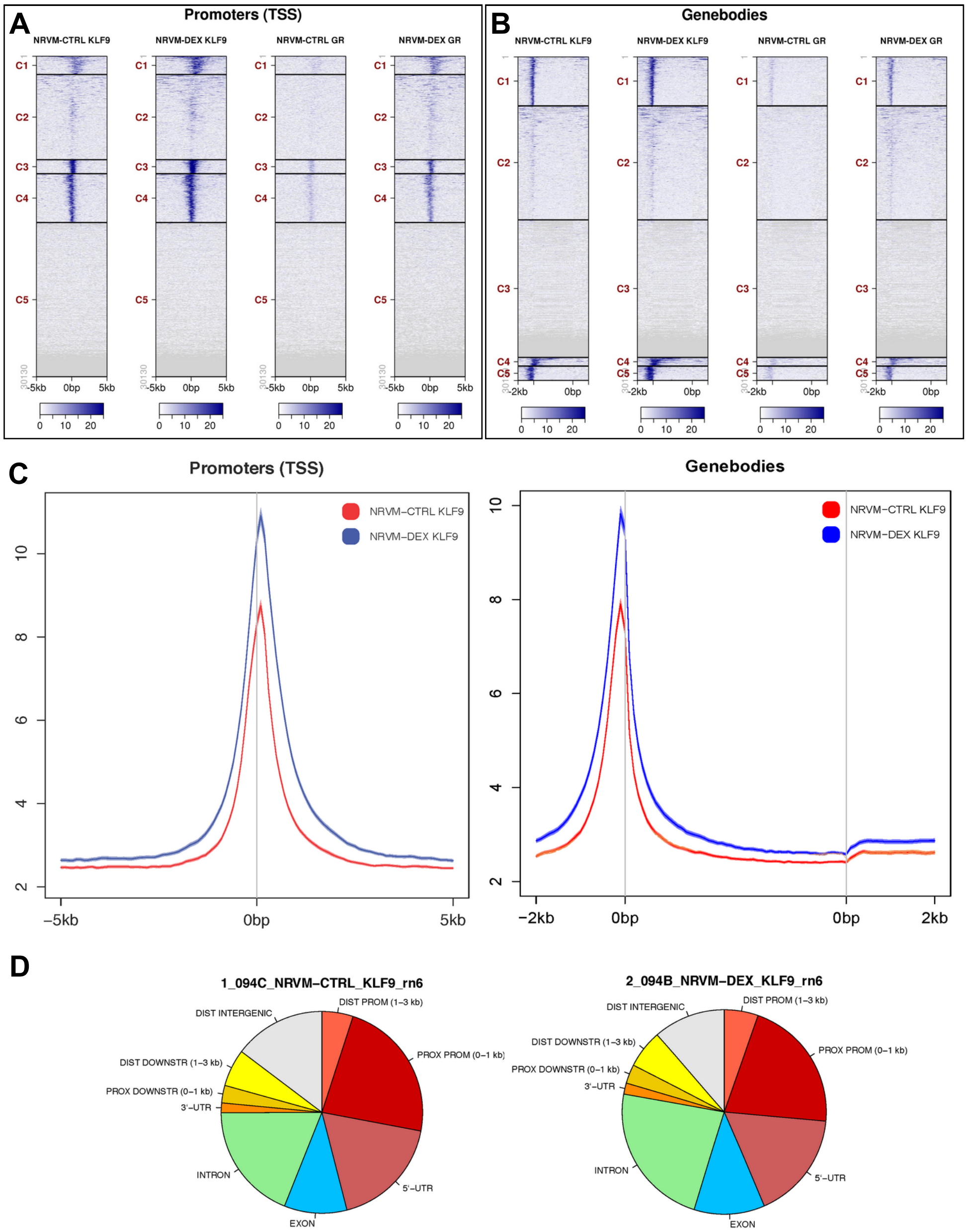


Figure S2. A and B. Heatmap showing tag distribution for Klf9 and GR across the active regions at the propoters and the gene bodies, (values in z-axis/color, active regions in y-axis) in control and Dex treated cardiomyocytes (6hrs for Klf9-ChIP-Seq and 1hr for GR-ChIP-Seq). The date is presented in 5 clusters (default), C1 to C5 and sorted. C. Average plots of Klf9 tag distribution of active regions (promoters and gene bodies) in control (red), Dex 6hrs treated (blue) cardiomyocytes. D. Pie chart showing the location of the Klf9 peaks relative to genomic annotations in control and Dex (100nM, 6hrs) treated cardiomyocytes.

### Figure S3

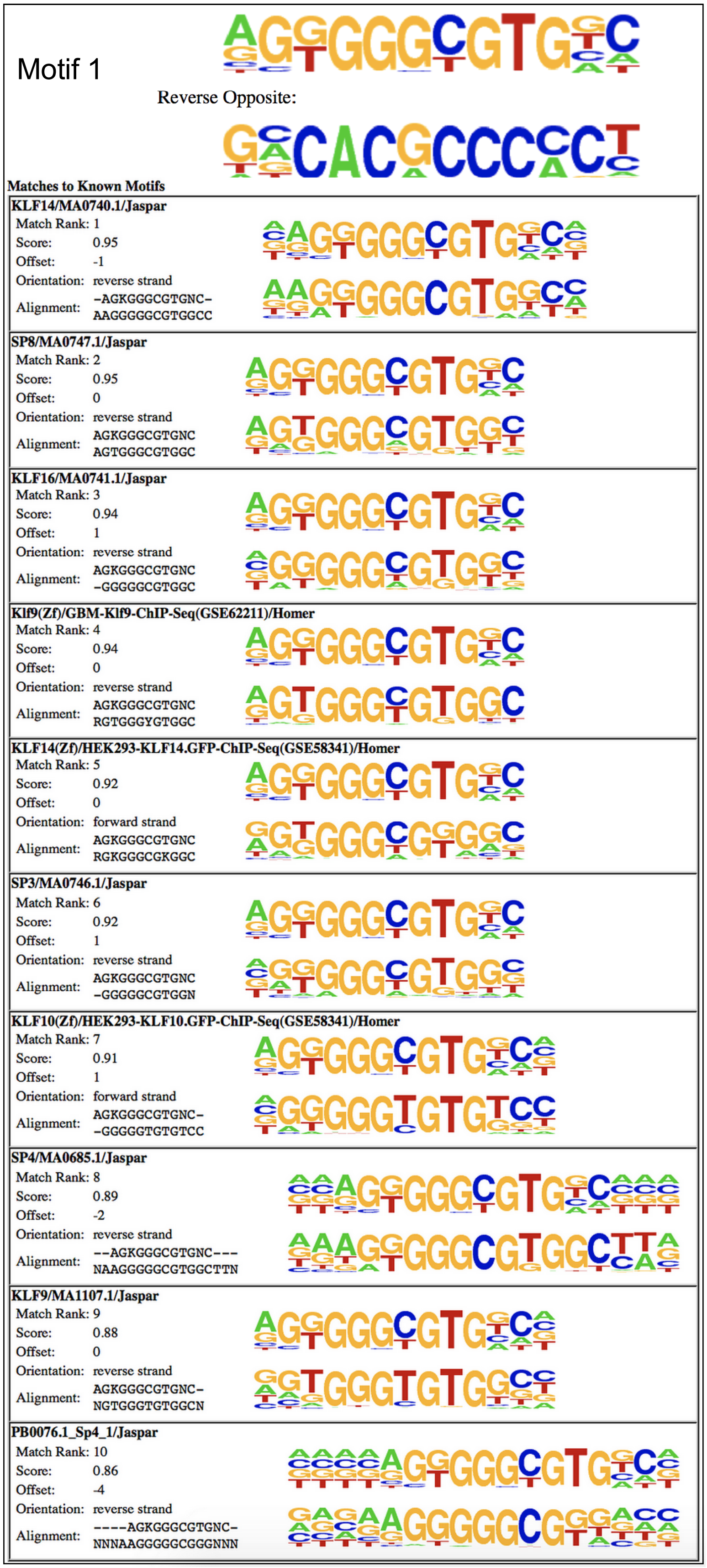


**Homer *de novo* Motif Results (QC/homer/2_094B_NRVM-DEX_KLF9_rn6/)**

Known Motif Enrichment Results
Total target sequences = 2491
Total background sequences = 39204

| Rank | Motif | P-value | log P-pvalue | % of Targets | % of Background |
|---|---|---|---|---|---|
| 1 | 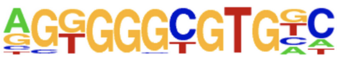 | 1e-142 | -3.291e+02 | 10.84% | 1.42% |
| 2 | 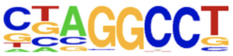 | 1e-123 | -2.853e+02 | 54.80% | 31.73% |
| 3 | 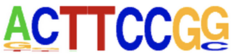 | 1e-86 | -1.994e+02 | 35.17% | 18.40% |
| 4 | 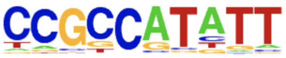 | 1e-48 | -1.116e+02 | 14.65% | 6.34% |
| 5 | 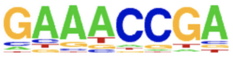 | 1e-35 | -8.239e+01 | 57.89% | 45.31% |
| 6 | 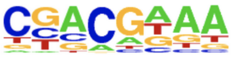 | 1e-32 | -7.555e+01 | 62.51% | 50.52% |
| 7 | 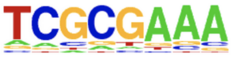 | 1e-31 | -7.239e+01 | 54.40% | 42.65% |
| 8 | 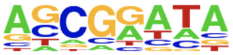 | 1e-28 | -6.463e+01 | 26.54% | 17.59% |
| 9 | 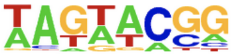 | 1e-26 | -6.075e+01 | 16.46% | 9.56% |
| 10 | 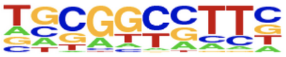 | 1e-21 | -4.935e+01 | 6.46% | 2.76% |

Figure S3. HOMER motif analysis. Regions of 200bp surrounding the summit of the top 2,500 peaks (based on MACS2 p-values) are analyzed. identified enriched de novo motifs are mathced with HOMER's internal motif database.
http://homer.ucsd.edu/homer/motif/

### Figure S4

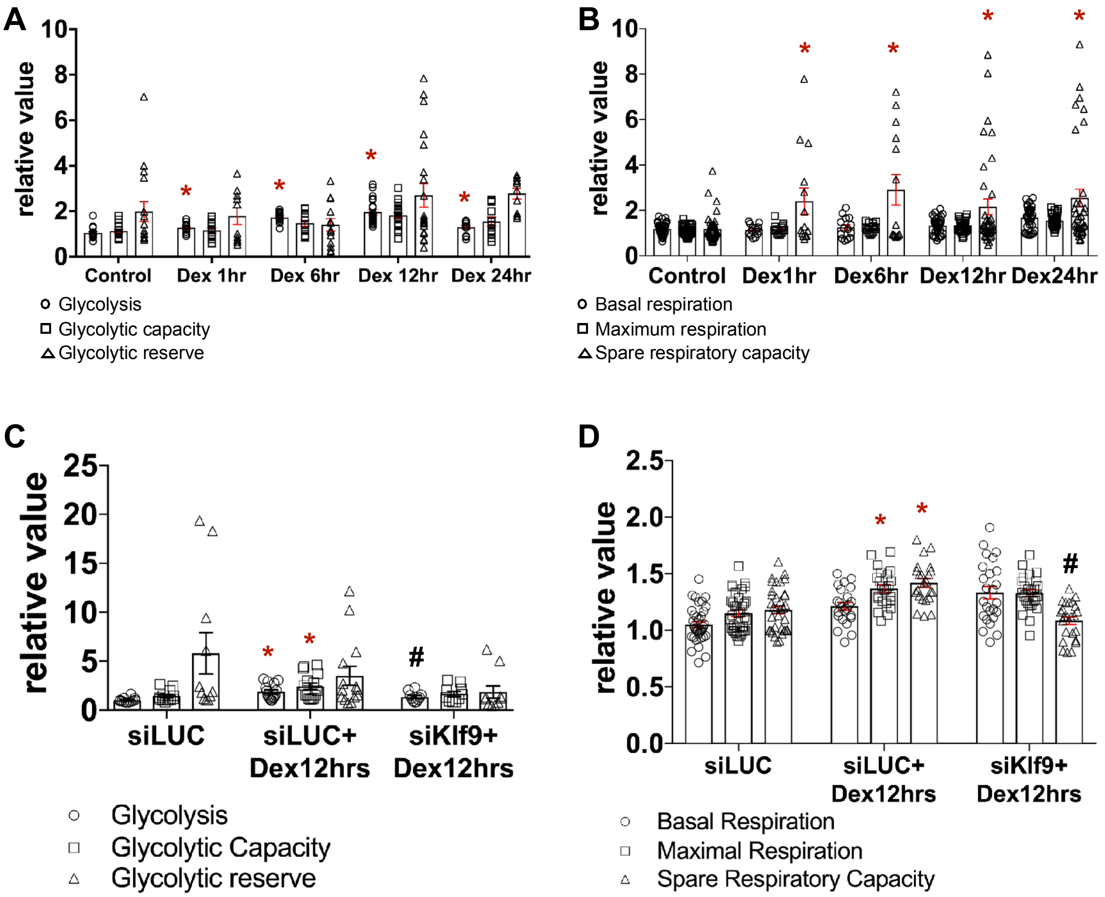


Figure S4. A. Graph represents Glycolysis, Glycolytic capacity and Glycolytic reserve in cardiomyocytes treated with control (methanol) or Dex (100nM) for increaseing time periods, as indicated, and measured by the Glucolysis stress test and calculated by report generator of the Sea horse software. Each point is data from individual well, from three indepdendent cardiomyocyte culture. Error bars represents SEM, and p-value is <0.5 compared to control. B. Graph represnts Basal respiration, Maximum respiration and Spare respirtory capacity, in cardiomyocytes treated with control (methanol) or Dex (100nM) for increaseing time periods, as indicated, and measured by the Mito stress test and calculated by report generator of the Sea horse software. Each point is data from individual well, from three indepdendent cardiomyocyte culture. Error bars represents SEM, and p-value is <0.5 compared to control. C. Graph represents Glycolysis, Glycolytic capacity and Glycolytic reserve in cardiomyocytes treated with control (methanol) or Dex (100nM) with or without ad-siKlf9 mediated Klf9 knockdown, as indicated, and measured by the Glucolysis stress test and calculated by report generator of the Sea horse software. Each point is data from individual well, from three indepdendent cardiomyocyte culture. Error bars represents SEM, and p-value is <0.5 compared to control. D. Graph represnts Basal respiration, Maximum respiration and Spare respirtory capacity, in cardiomyocytes treated with control (methanol) or Dex (100nM) with or without ad-siKlf9 mediated Klf9 knockdown as indicated, and measured by the Mito stress test and calculated by report generator of the Sea horse software. Each point is data from individual well, from three indepdendent cardiomyocyte culture. Error bars represents SEM, and p-value is <0.5 compared to control.

Antibody list

| Antibody | Company | Cat# |
|---|---|---|
| Nr3c1 (GR) | Cell Signalling Technology | 3660S |
| Ankrd1 (CARP) | Santacruz | sc-365056 |
| Gapdh | Cell Signalling Technology | 97166S |
| H2b | Cell Signalling Technology | 12364S |
| Klf9 | Abclonal | A7196 |
| | Invitrogen | 701888 |
| | Origene | TA324593 |
| Klf15 | Abclonal | A7194 |
| | Sigma | AV32587 |

TaqMan qPCR Assays

| Gene | ID |
|---|---|
| Arrdc3 | Rn01757892_m1 |
| Hes1 | Mm01342805_m1 |
| Klf10 | Mm00449812_m1 |
| Klf15 | Mm00517792_m1 |
| Klf2 | Rn01420496_gh |
| Klf3 | Rn01413619_m1 |
| Klf6 | Mm00516184_m1 |
| Klf9 | Mm00495172_m1 |
| Myh6 | Mm00440359_m1 |
| Myh7 | Mm00600555_m1 |
| Nppa | Mm01255747_g1 |
| Nr3c1 | Rn00561369_m1 |
